# Exploiting the Nonlinear Structure of the Antithetic Integral Controller to Enhance Dynamic Performance

**DOI:** 10.1101/2022.08.02.502513

**Authors:** Maurice Filo, Sant Kumar, Stanislav Anastassov, Mustafa Khammash

## Abstract

The design of biomolecular feedback controllers has been identified as an important goal across a broad range of biological applications spanning synthetic biology, cell therapy, metabolic engineering, etc. This originates from the need to regulate various cellular processes in a robust and timely fashion. Recently, antithetic integral controllers found their way into synthetic biology due to the Robust Perfect Adaptation (RPA) property they endow — the biological analogue of robust steady-state tracking. The antithetic integral motif hinges on a sequestration reaction between two molecules that annihilates their function. Here, we demonstrate that the complex resulting from the nonlinear sequestration reaction can be leveraged as an inhibitor to enhance the dynamic performance while maintaining the RPA property. We establish that this additional inhibition by the sequestration complex gives rise to a filtered Proportional-Integral (PI) controller thus offering more flexibility in shaping the dynamic response and reducing cell-to-cell variability. Furthermore, we explore the effect of various biological inhibitory mechanisms on the overall performance. The various analyses in the paper are carried out using analytical tools and are supported by numerical simulations. Finally, an experimental validation is performed using the cyberloop — a hybrid platform where the controller is implemented *in silico* to control a genetic circuit *in vivo*.

## I. INTRODUCTION

One of the essential features of living systems is their ability to maintain a robust behavior despite disturbances coming from their external uncertain and noisy environments. This feature, in pure biological terms, is referred to as homeostasis which is typically achieved via endogenous feedback regulatory mechanisms shaped by billions of years of evolution inside the cells. Pathological diseases are often linked to loss of homeostasis [1], [2]. The need to restore homeostasis for preventing such diseases, when endogenous regulatory mechanisms fail, was one of the main drivers of ushering a new active field of research referred to as cybergenetics [3] — a field that brings control theory and synthetic biology together. In particular, the rational design of biomolecular feedback controllers offers promising candidates that may accompany or even replace such failed mechanisms [4]–[6].

A notion which is similar to homeostasis, but more stringent, is Robust Perfect Adaptation (RPA) (see e.g. [7], [8]) which is the biological analogue of the well-known notion of robust steady-state tracking in control theory. A controller succeeds in achieving RPA if it drives the steady state of a variable of interest to a prescribed set-point despite varying initial conditions, plant uncertainties and/or constant disturbances. Motivated by the internal model principle [9], which establishes that the designed controller must implement an integral feedback component to be able to achieve RPA, the antithetic integral controller [10] was brought forward. The basic antithetic integral motif interconnected in a feedback configuration with an arbitrary network (Figure 1(a)) is depicted in Figure 1(b). To see the RPA property endowed by this motif, one can write the controller dynamics as

**Fig. 1.**
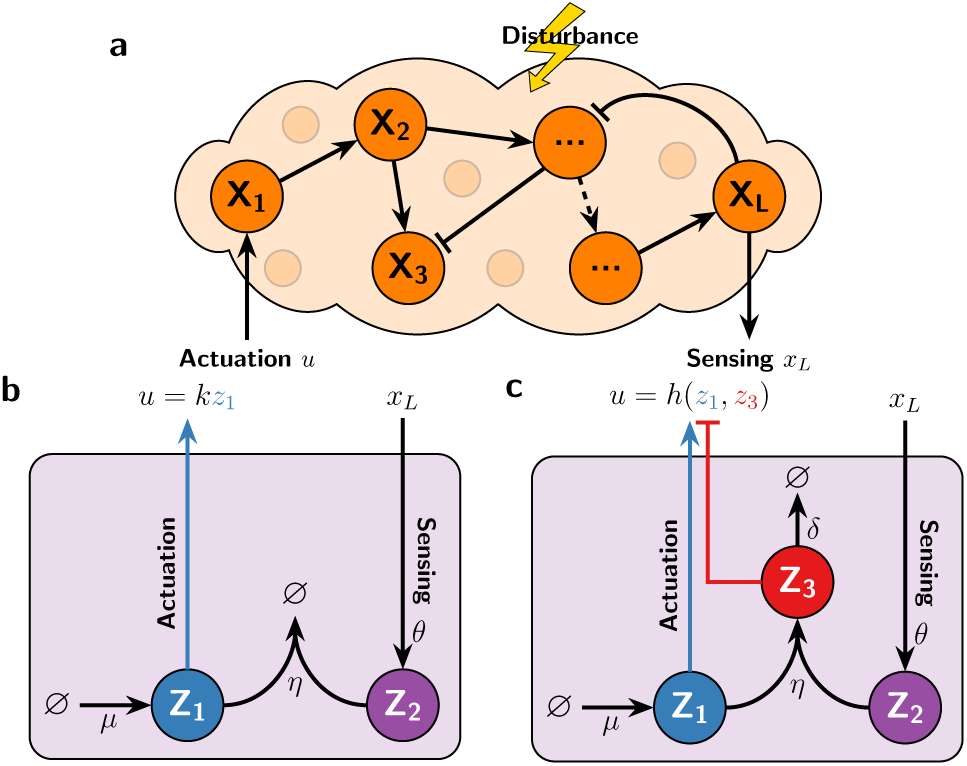
Antithetic Integral Feedback Motif. **(a)** The plant: an arbitrary network to be controlled. The input and output of this network are denoted by **X**_**1**_ and **X**_**L**_, respectively. The feedback controller senses the output **X**_**L**_ and actuates the input **X**_**1**_ accordingly. **(b)** The basic antithetic integral motif. The controller molecule **Z**_**1**_ is constitutively produced at a rate *µ*, while **Z**_**2**_ is catalytically produced by the output **X**_**L**_ at a rate *θx*_*L*_. Furthermore, **Z**_**1**_ and **Z**_**2**_ sequester each other at a rate *η* to produce an inert complex ∅ that has no effect on the closed-loop dynamics. **(c)** Exploiting the sequestration complex as an inhibitor. The sequestration reaction produces a non-inert molecule denoted by **Z**_**3**_ that participates, together with **Z**_**1**_, in the actuation of the plant. **Z**_**1**_ acts as an activator while **Z**_**3**_ acts as an inhibitor. Note that *δ* denotes the removal rate of the sequestration complex **Z**_**3**_.

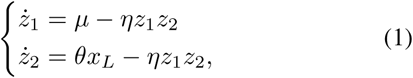

where lowercase letters denote the concentrations of the associated species represented by uppercase bold letters. Assuming closed-loop stability, the steady-state concentration of **X**_**L**_, denoted by 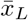, is *µ/θ* and is independent of the plant parameters, disturbances and initial conditions. Hence RPA will be ensured as long as the closed-loop network is stable. In fact, it is shown in [11] that, for stochastic networks, the antithetic integral motif is necessary to achieve RPA. Ever since the antithetic integral controller was introduced, more focus was directed on enhancing its performance either by tuning and exploring the dynamic trade-offs [12]–[15], or by appending additional circuitry [16]–[21] including molecular buffering and realizations of PI and PID controllers. Furthermore, biological implementations of such integral controllers [11], [22]–[26] have recently received more focused attention.

At the heart of the antithetic motif lies the nonlinear sequestration reaction between the controller molecules **Z**_**1**_ and **Z**_**2**_ which produces an inert complex ∅. In this work, we instead propose a simple idea to leverage this nonlinearity by exploiting the sequestration complex, denoted by **Z**_**3**_, as an inhibitor (see Figure 1(c)) to enhance the overall dynamic performance of the antithetic integral controller. We first establish, in Section II, that this additional inhibition augments the integrator with a filtered proportional component to yield a filtered PI controller. Subsequently, we derive the mappings that link the PI gains and cutoff frequency of the filter to the biomolecular parameters for various biologically-relevant inhibitory mechanisms in Section III. These mappings aid in the tuning of the molecular controllers and uncover their different degrees of flexibility. In sections IV and V, we carry out an analysis, backed up with simulations, to demonstrate the flexibility brought by the inhibitory sequestration complex in shaping the deterministic dynamics and reducing the variance in the stochastic setting. Finally before concluding, we provide an experimental validation of our biomolecular controller in Section VII using a recently developed hybrid platform — the cyberloop [26], [27] — where the controller is implemented *in silico* to regulate a biological genetic circuit in yeast.

## II. Unravelling the Controller Structure

### A. Controller Descriptionxs

Consider the molecular controller proposed in Figure 1(c), interconnected in feedback with an arbitrary plant comprised of *L* species: **X**_**1**_, **X**_**2**_, …, **X**_**L**_. The difference between this controller and the basic antithetic integral motif in Figure 1(b) is that the sequestration reaction between the controller molecules **Z**_**1**_ and **Z**_**2**_ produces a non-inert complex, denoted by **Z**_**3**_, which is capable of inhibiting the plant. To study different inhibition mechanisms, the controller is allowed to actuate the input species **X**_**1**_ of the plant via a production reaction and a degradation reaction given by

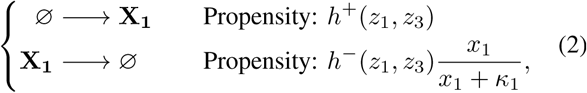

where the functional forms of *h*^*±*^ dictate the specific inhibition mechanism such as degradation, competitive or additive inhibition (see Section III for more details). Consequently, by defining the total control action, denoted by *u*, as

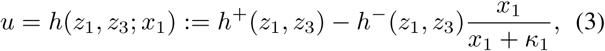

we can write the general closed-loop dynamics, for any inhibition mechanism, as

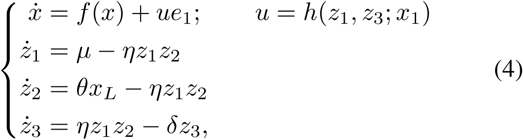

where *x* := (*x*_1_ *x*_2_ *… x*_*L*_]^*T*^, and *e*_*i*_ denotes the vector with a one in the *i*^th^ coordinate and zeros elsewhere. Note that *h* is chosen to be a monotonically increasing function of *z*_1_ and a decreasing function in *z*_3_ to capture the activation and inhibition mechanisms by **Z**_**1**_ and **Z**_**3**_, respectively.

### B. Linear Perturbation Analysis

In this section, we uncover the inhibitory effect of the sequestration complex **Z**_**3**_ on the control architecture using linear perturbation theory. In fact, we show that the overall control structure embeds a filtered PI controller.

Let 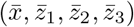 denote the fixed point of the closed-loop network operating at a nominal exogenous input 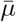 which prescribes the output set-point to 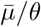. Furthermore, let 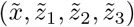 and 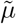 denote a perturbation from the closed-loop fixed point and nominal exogenous input, respectively. That is, we have

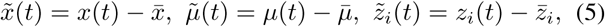

for *i* = 1, 2, 3. The linearized dynamics are thus given by

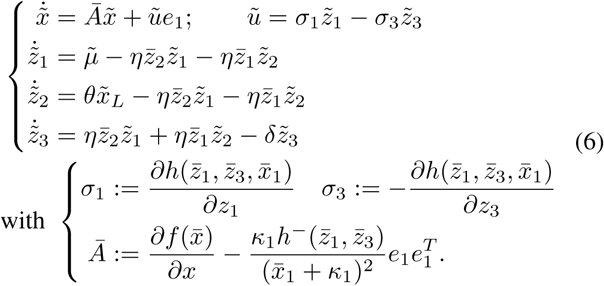

Observe that due to the monotonicity of *h*, we have *σ*_1_, *σ*_3_ *>* 0. The underlying control structure is most easily uncovered and visualized by drawing the block diagram of the linearized dynamics (refer to [16]). Carrying out some algebraic manipulations in the Laplace domain, one can write the Laplace transform of the control action, denoted by *ũ* (*s*), in terms of the Laplace transforms of the exogenous input 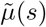, output 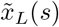 and error 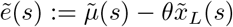 as

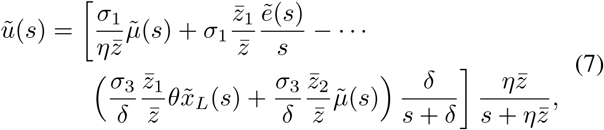

where 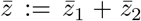. The filtered PI structure can already be seen from (7); however, this structure is most easily seen in the strong sequestration regime (large *η*). Assuming that the set-point is admissible (i.e. there exists a positive input *u* that achieves the desired set-point), it can be shown that 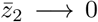 as *η* → ∞, and therefore the controller transfer function in (7) simplifies to

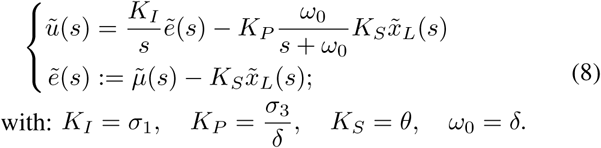

Equipped with the controller and plant transfer functions given by (8) and 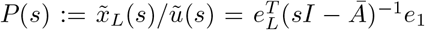, respectively, we can now draw the block diagram shown in Figure 2 to reveal the underlying filtered PI-control structure.

**Fig. 2.**
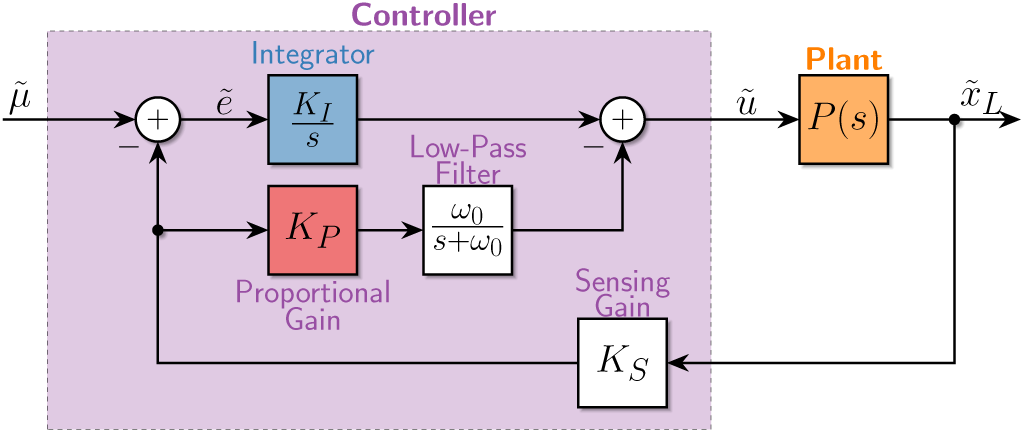
Block Diagram for the linearized closed-loop dynamics of (4). The block diagram reveals that the underlying control architecture involves an integral controller with gain *K*_*I*_ and a filtered-proportional controller with gain *K*_*P*_ and cuttoff frequency *ω*_0_ given in (8). The integral component acts on the error signal 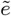 which is necessary to achieve RPA; whereas, the filtered proportional component acts on the output 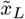 directly (see [28, Chapter 10] for more details on this class of controllers that act on the error and the output separately and simultaneously).

## III. Mappings between the Filtered-PI and Biomolecular Parameter Spaces

Next, we derive the mappings, in the strong-sequestration regime, between the filtered-PI parameters (*K*_*P*_, *K*_*I*_, *ω*_0_) and the various biomolecular parameters (*θ, δ*, …) using a similar approach as that in [16]. We first start with the analysis problem: given the biomolecular parameters, what are the PI gains and cutoff frequency? Then we move to the design problem: what are the biomolecular parameters that achieve some desired PI gains and cutoff frequency?

Throughout the subsequent analysis, we will make two assumptions about the plant. Let *F*_*i*_ (*i* = 1, 2, …, *L*) denote the steady-state maps of the plant, that is, if *u* is a constant then with reference to the first equation in (4), we write

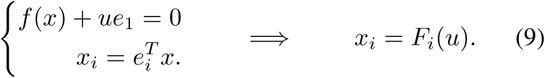

### Assumption 1

Assume that for the desired steady-state output 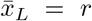, there exists a supporting input, denoted by *ū*, that achieves the desired output. More precisely, for *r >* 0, ∃*ū*∈ 𝒰⊂ ℝ such that *F*_*L*_(*ū*) = *r*, where 𝒰 represents the set of admissible inputs to the plant.

### Assumption 2

Assume that for the admissible supporting input *ū*, the steady-state concentrations of the internal plant species, denoted by 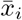, *i* = 1, 2, *…, L* − 1, exist and are non-negative. More precisely, for 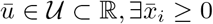 such that 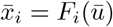 for *i* = 1, 2,, *L* − 1.

We emphasize that these assumptions do not depend on the type of controller used. Instead, they only depend on the plant and the particular choice of actuated input species. These assumptions have to be satisfied, otherwise the choice of the input species is simply inadequate and there is no controller that can achieve the desired output without changing the choice of the actuated input species. For simplicity, we let 𝒰= ℝ_+_; however, this can be relaxed to include negative inputs *u* as well (that can be achieved with degradation inhibition for example), but with a lower bound that depends on the particular mechanism *h* at hand. Next, we treat the analysis and design problems for three biologically-relevant actuation functions *h* with different inhibition mechanisms: additive, competitive and degradation.

### A. Additive Inhibition

Consider an arbitrary plant, satisfying Assumptions 1 and 2, regulated by the filtered-PI controller given in (4) where **Z**_**3**_ separately produces the input **X**_**1**_ in a repressive manner. The actuation function *h* is thus given by

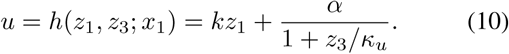

The set-point is given by 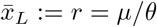. For a given plant and set-point *r*, the supporting input *ū* satisfies *F*_*L*_(*ū*) = *r* and is fixed (see Assumption 1). We first treat the analysis problem, then move on to the design problem.

#### a) Analysis

The controller coordinates 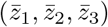 of the fixed point are given by

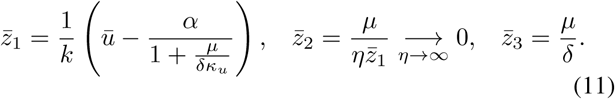

Clearly, the following condition on the biomolecular parameters has to be satisified to guarantee that 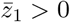,

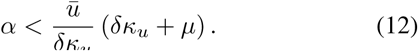

Violating this condition causes one of the coordinates of the fixed point to become negative and thus causing instability. Using the formulas derived in (8) and the partial derivatives defined in (6), one can write the PI gains (*K*_*P*_, *K*_*I*_) and cutoff frequency *ω*_0_ in terms of the various biochemical parameters as

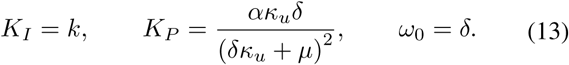

Observe that the gains and cutoff frequency are all independent of the plant.

#### b) Design

By fixing *κ*_*u*_, *µ* and *r* (and thus *ū*), one can easily solve the equations given in (13) for the biomolecular parameters *k, α* and *δ* in terms of the PI gains and cutoff frequency to obtain

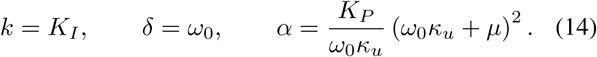

Since the biomolecular parameters *k, α* and *δ* cannot be negative and have to satisfy the condition in (12), the achievable PI gains and cutoff frequency are constrained as described next.

#### c) Filtered-PI Coverage

Constraining *k, α* and *δ* to be non-negative and to satisfy condition (12) yields the following achievable PI gains and cutoff frequency.

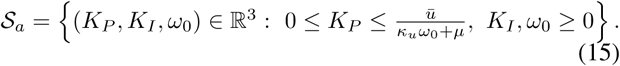

This means that with additive inhibition, the proportional gain *K*_*P*_ has an upper bound. Values of *K*_*P*_ larger than the upper bound cannot be achieved without causing a negative equilibrium (and thus losing stability), no matter how the biomolecular parameters are tuned. In fact, the highest achievable *K*_*P*_ is precisely the bifurcation point. Finally, note that this upper bound depends on the plant and desired set-point via the supporting input *ū*.

### B. Competitive Inhibition

Next, consider the case where **Z**_**3**_ competitively inhibits the input **X**_**1**_. The actuation function *h* is thus given by

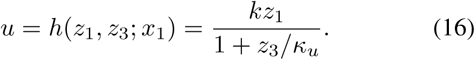

The controller coordinates of the fixed point are given by

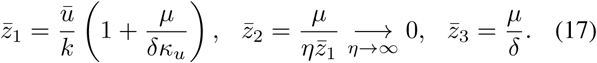

Clearly, 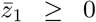 when *ū ∈* 𝒰 = ℝ_+_. Carrying out calculations similar to the additive inhibition case, we obtain the following parameter mappings

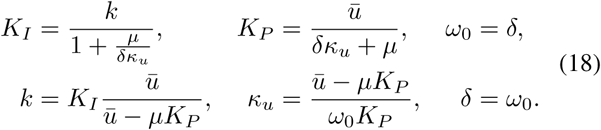

Constraining *k, κ*_*u*_ and *δ* to be non-negative yields the following achievable PI gains and cutoff frequency.

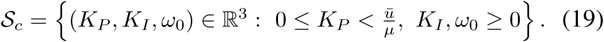

This means that with competitive inhibition, similar to the additive inhibition, the proportional gain *K*_*P*_ has an upper bound. Values of *K*_*P*_ larger than the upper bound cannot be achieved no matter how the biomolecular parameters are tuned. However, unlike additive inhibition, there is no risk of negative equilibrium caused by the controller. Furthermore, observe that *κ*_*u*_ tunes both the proportional and integral gain simultaneously and in an opposite direction: decreasing *κ*_*u*_ increases the proportional gain while decreasing the integral gain. This has an attractive feature that favors stability of the closed-loop dynamics.

### C. Degradation Inhibition

Next, consider the case where **Z**_**3**_ degrades the input **X**_**1**_. The actuation function *h* is thus given by

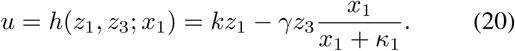

The set-point is given by 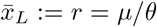. For a given plant and set-point *r*, the supporting input *ū* and input species concentration 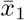 satisfy *F*_*L*_(*ū*) = *r* and 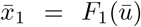, respectively, and are both fixed (see Assumptions 1 and 2). The controller coordinates of the fixed point are given by

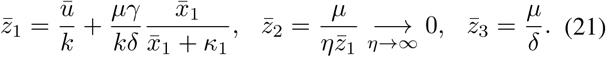

Clearly, 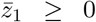 when *ū* ∈ 𝒰= ℝ_+_. Carrying out calculations similar to the additive inhibition case, we obtain the following parameter mappings

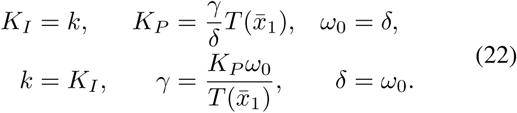

where 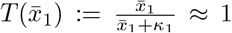, by choosing *κ*_1_ to be small enough. Note that this is not a requirement for the controller; in fact, *κ*_1_ can be used as an additional tuning parameter that can be shown to have benefits on shaping the dynamic response. This stems from the fact that, for nonzero *κ*_1_, the controller embeds two proportional feedback actions from *x*_1_ (state feedback) and *x*_*L*_. However, our focus here is to track the effects of the output feedback alone to make a fair comparison with the other inhibition mechanisms (refer to the Supplementary Information in [16] for more details). Constraining *k, γ* and *δ* to be non-negative yields the following achievable PI gains and cutoff frequency.

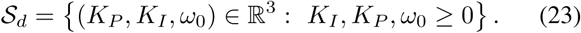

This means that with degradation inhibition, the filtered-PI controller can achieve all positive gains and cutoff frequency by suitably tuning the biomolecular parameters *k, γ* and *δ*.

By inspecting the coverages given in (15), (19) and (23), one can clearly see that 𝒮_*a*_ ⊂ 𝒮_*c*_ ⊂ 𝒮_*d*_ as visually demonstrated in Figure 3. In conclusion, this analysis demonstrates that degradation inhibition offers more tuning flexibility than the competitive inhibition which in turn offers more tuning flexibility than additive inhibition.

**Fig. 3.**
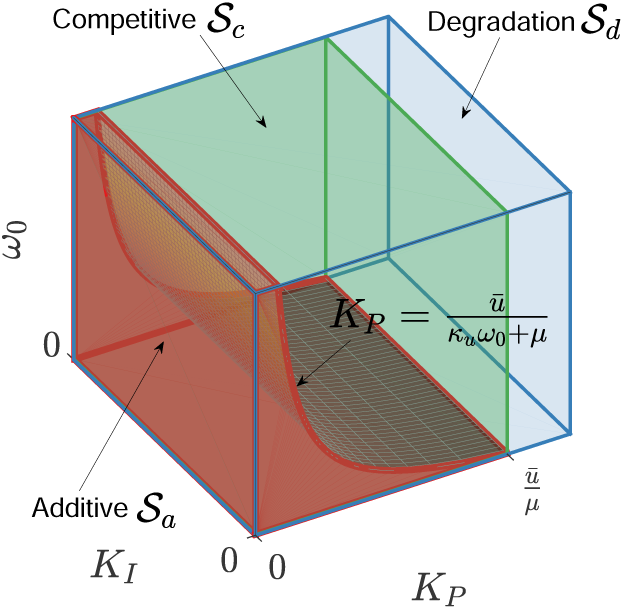
Filtered-PI Coverage. The colored regions show the PI gains (*K*_*P*_, *K*_*I*_) and cutoff frequency *ω*_0_ that are achievable by tuning suitable biomolecular parameters. The regions are color coded for different actuation functions *h* modeling the three different inhibition mechanisms: additive (10), competitive (16) and degradation (20). Note that *ū* is the steady-state supporting input required to achieve the desired set-point, and its value depends only on the plant and the desired set-point. The spans of achievable filtered-PI parameters for the cases of additive, competitive and degradation inhibitions are respectively calculated as 𝒮_*a*_ in (15), 𝒮_*c*_ in (19) and 𝒮_*d*_ in (23) and shown here to satisfy 𝒮_*a*_ ⊂ 𝒮_*c*_ ⊂ 𝒮_*d*_. This illustrates that degradation offers more tuning flexibility than competitive inhibition which in turn offers more tuning flexibility than additive inhibition.

## IV. Dynamic Performance Assessment

The objective of this section is to demonstrate analytically and through simulations, that designing **Z**_**3**_ to have an inhibitory effect on the input **X**_**1**_ (i.e. filtered-PI control), enables more flexibility in enhancing the dynamic performance when compared to the case where **Z**_**3**_ is inert (i.e. standalone antithetic integral motif). We also explore the performance-enhancement capabilities of the different inhibition mechanisms presented in Section III.

Consider the closed-loop dynamics given in (4), where the control action *u* can be any of the three mechanisms involving degradation, competitive or additive inhibition. Using the block diagram in Figure 2, it is straightforward to write down the closed-loop transfer functions of the linearized dynamics from 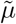 to 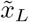 for the I (*K*_*P*_ = 0) and filtered-PI (*K*_*P*_ *>* 0) cases

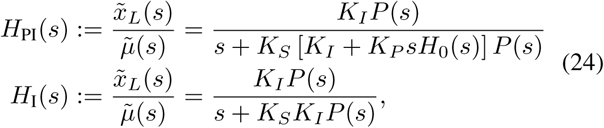

where 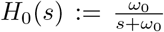 is the transfer function of the low-pass filter. For simplicity, we consider the plant to be a simple birth-death process with *L* = 1 species depicted in Figure 4(a). That is *f* (*x*) = − *γ*_1_*x* + *u* in (4). This simple plant is enough to demonstrate the added flexibility. The transfer function of this plant is given by 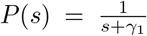. By substituting for *P* (*s*) in *H*_I_(*s*), we obtain

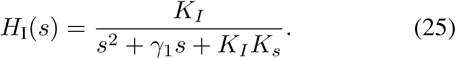

**Fig. 4.**
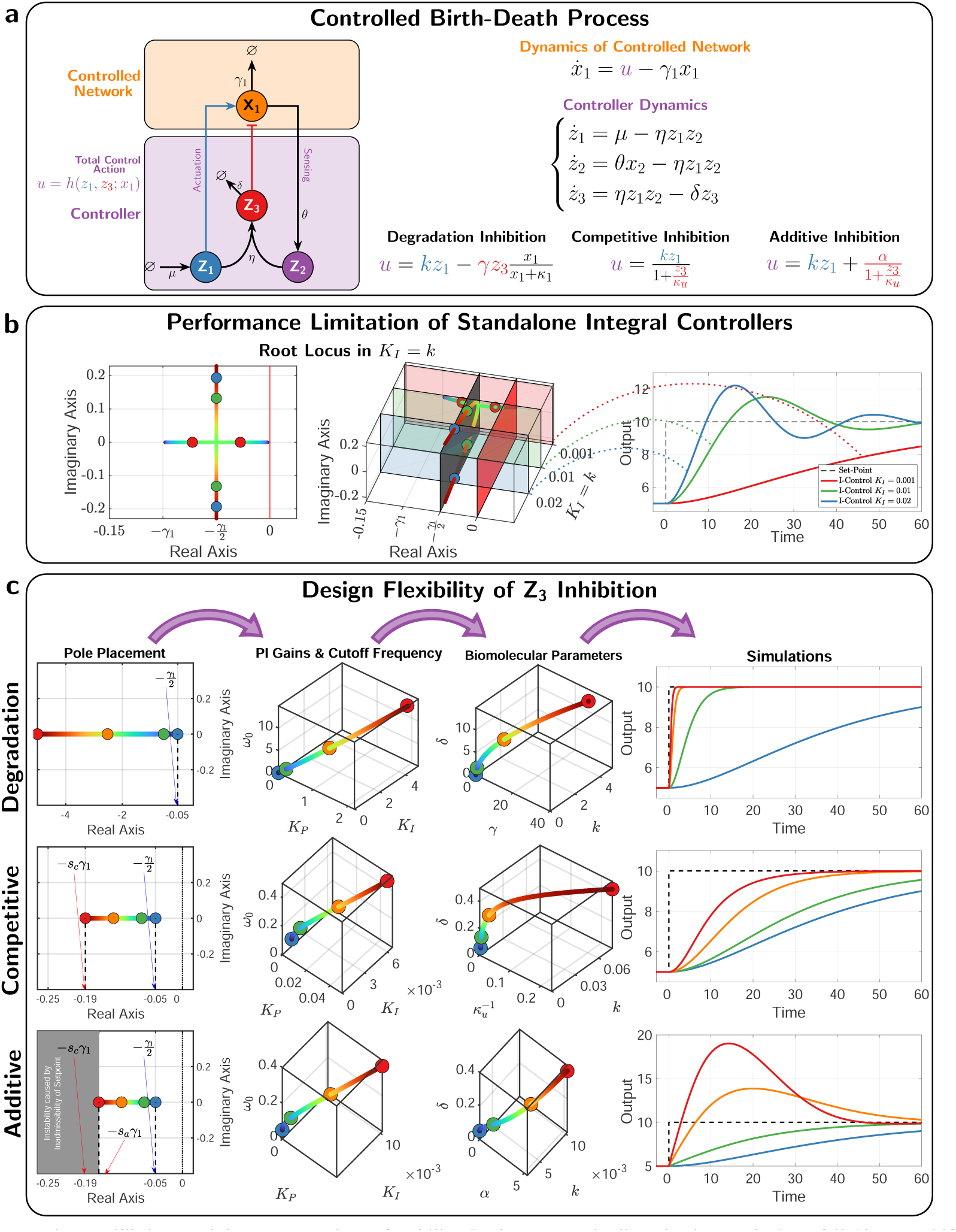
Performance Flexibility Offered by an Inhibitory Sequestration Complex Z_3_. (a) Closed-loop dynamics. A birth-death process is controlled by the antithetic integral controller that exploits the sequestration complex as an inhibitor. The control action *u* is shown here for three types of inhibition mechanisms. (b) Performance limitation of the standalone antithetic integral controller. Without the inhibiting sequestration complex **Z**_**3**_ (i.e. *u* = *kz*_1_), the response cannot be sped up beyond a certain threshold without inflicting oscillations. The left and middle plots demonstrate the same root locus of the linearized dynamics as the integral gain *K*_*I*_ is increased. The left plot depicts the complex plane, while the middle plot explicitly shows the complex plane together with the values of the integral gain *K*_*I*_ which is shown to be approximately equal to *k*. These plots verify that two eigen-values are confined within a small region close to the imaginary axis when *γ*_1_ is small, and thus imposing a limitation on the achievable performance as demonstrated in the simulations shown in the right plot. (c) Design flexibility offered by the inhibitory sequestration complex. Giving rise to a filtered-PI controller, the inhibitory sequestration complex offers more flexibility in achieving superior performance compared to the standalone antithetic motif. This panel shows the steps of a pole-placement control design problem where the three dominant poles are placed on the real axis of the left-half plane to ensure a stable and non-oscillating response with a minimal overshoot. With competitive and additive inhibitions, there is a restriction on how far to the left the poles can be placed; whereas with degradation inhibition, one can place the poles arbitrarily as far to the left as desired and thus achieving a response that is as fast as desired without overshoots nor oscillations. However, with additive inhibition, placing the poles far to the left gives rise to a negative equilibrium and thus causes a loss of stability. It also causes the linearization analysis to fail (due to a bifurcation) as demonstrated by the large overshoots in the nonlinear simulations that are not predicted by the linearized analysis. The design problems start by picking the poles, then computing the PI gains and cutoff frequency, and finally computing the actual biomolecular parameters that allows us to obtain the nonlinear simulations to the right. The numerical values of the common parameters are *γ*_1_ = 0.1, *µ* = 5, *θ* = 1, *η* = 10. For degradation inhibition, *κ*_1_ = 0.1 and (*k, γ, δ*) are tuned. For competitive inhibition, (*k, κ*_*u*_, *δ*) are tuned. For additive inhibition, *κ*_*u*_ = 5 and (*k, α, δ*) are tuned. To change the set-point at *t* = 0, *µ* is doubled.

The poles of *H*_1_ (*s*) are given by 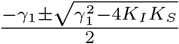. This explicitly provides the root locus of the closed-loop that is shown in Figure 4(b). The two poles start at *s* = 0 and *s* = − *γ*_1_ for *K*_*I*_ = 0, and as *K*_*I*_ is increased the two poles move on the real line closer to each other until they merge for 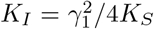. Increasing *K*_*I*_ further gives rise to two complex conjugate poles moving on the line *s* = −*γ*_1_*/*2 and diverging to −*γ*_1_*/*2 *±i*∞ as *K*_*I*_ → ∞. Observe that no matter how we tune *K*_*I*_, the poles cannot be placed to the left of *s* = −*γ*_1_. This means that the performance of the integral controller alone is limited, in the sense that the speed of the response cannot exceed a threshold dictated by *γ*_1_ without incurring overshoots and oscillations as demonstrated in Figure 4(b).

This is exactly where the filtered-proportional component, brought by the inhibitory nature of **Z**_**3**_, comes into play to add more flexibility. By substituting for *P* (*s*) in *H*_PI_(*s*), we obtain

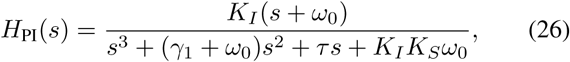

where *τ* := *γ*_1_*ω*_0_ + *K*_*I*_*K*_*S*_ + *K*_*P*_ *K*_*S*_*ω*_0_. To demonstrate the added flexibility, consider the pole placement design problem, where one aims at designing the values of the PI gains (*K*_*P*_, *K*_*I*_) and the cutoff frequency *ω*_0_ to place the three closed-loop poles at *s* = −*a*. Ideally, we aim at placing the poles in the left-half complex plane to guarantee stability and on the real axis to avoid oscillations. Furthermore, to obtain a fast response, the poles should be placed far to the left in the left-half complex plane. These criteria can all be achieved if we make *a >* 0 and large enough. Next, we study if this is possible to achieve with the filtered-PI controller using the three different actuation functions *u* presented in Figure 4(a). Note that for the standalone integrator, the only location that the two poles can be simultaneously placed at is *s* = − *γ*_1_*/*2 by setting 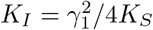. We show next that the added proportional component gives more flexibility. Since the three closed-loop poles are placed at *s* = − *a*, then the characteristic polynomial is given by

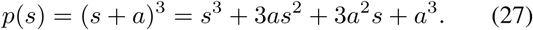

Equating *p*(*s*) to the denominator of *H*_PI_(*s*) allows us to express the designed PI gains (*K*_*P*_, *K*_*I*_) and cutoff frequency *ω*_0_ in terms of the birth-death parameter *γ*_1_, the sensing gain *K*_*S*_ and the placed pole −*a* as

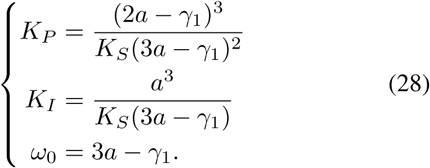

Recall, that the sets of achievable PI gains and cutoff frequencies for the three different inhibition mechanisms are given by 𝒮_*a*_, 𝒮_*c*_ and 𝒮_*d*_ in (15), (19) and (23). With some algebraic calculations (see Appendix A), it is easy to show that these sets constrain the achievable poles *s* = − *a* to the following regions on the real axis.

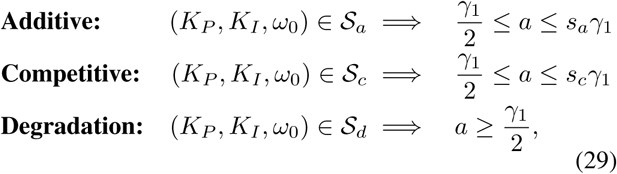

where 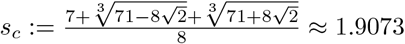 solves a cubic polynomial and *s*_*a*_ ∈ [*γ*_1_*/*2, *s*_*c*_] is a function of *κ*_*u*_*γ*_1_*/µ* and solves a quartic polynomial. As a result, with additive and competitive inhibitions, the fastest poles achievable are for *a* = *s*_*a*_*γ*_1_ and *a* ≈ 1.9073*γ*_1_, respectively, with *s*_*a*_ ≤ 1.9073. Whereas with degradation inhibition, there is no theoretical upper limit.

In conclusion, this case study (controlling the birth-death process) shows that the filtered-PI controller significantly improves the performance compared to standalone antithetic integral controllers. In particular, implementing the filtered-PI controller with a degradation inhibition enables us to make the response as fast as desired without giving rise to any overshoots or oscillations. However, although the other inhibition mechanisms also significantly improve the performance compared to the standalone antithetic integral controller, the responses cannot be made arbitrarily fast without incurring oscillations and/or overshoots. This is due to the reduced filtered-PI coverage (see (15) and (19)).

## V. Variance Reduction in the Stochastic Setting

It has been previously shown that proportional controllers are capable of reducing the output stationary (steady-state) variance in the stochastic setting [16], [17]. Here we show that using the sequestration complex as an inhibitor is also capable of reducing the output stationary variance as well. In addition to simulations, we carry out a tailored moment-closure technique similar to the one developed in [16], [17] in order to derive an approximate closed-formula for the stationary variance. The derivations, for the three inhibitory mechanisms, can be found in Appendix B. The results are summarized in Figure 5, where the plant is considered here to be a gene expression model such that **X**_**1**_ (mRNA) catalytically produces **X**_**2**_ (protein) at a rate *k*_1_ while both **X**_**1**_ and **X**_**2**_ degrade at a rate *γ*_1_ and *γ*_2_, respectively. Clearly, compared to the standalone antithetic integral controller 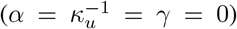, the various inhibitory mechanisms by the sequestration complex are all capable of reducing the output stationary variance. Observe that the approximate formula is capable of capturing the monotonicity of the stationary variance for all inhibition mechanisms. However, the accuracy of the approximate formula becomes worse as the parameters *α*, 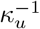 and *γ* (for the associated inhibitory mechanism) are increased. This is due to the non-linearity that kicks in more and thus making our tailored moment-closure technique less accurate.

**Fig. 5.**
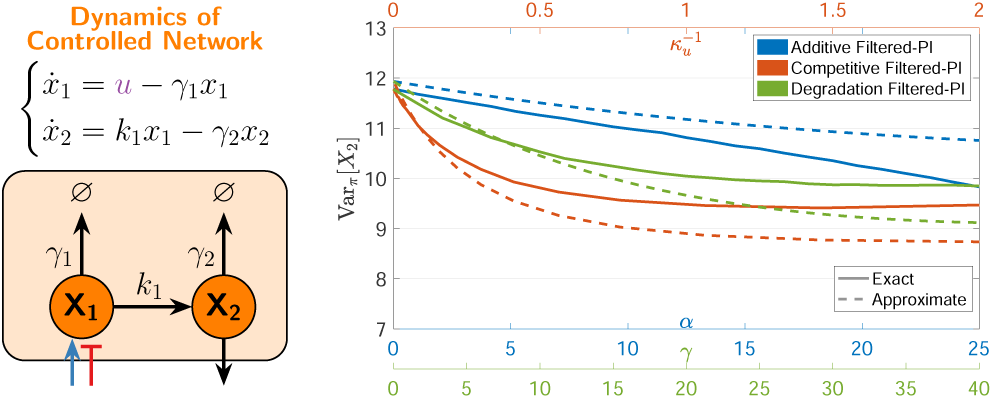
Variance Reduction by the Sequestration Complex. The plant considered here is a gene expression model depicted in the network to the left. The controller is the same as that presented in Figure 4(a). For additive, competitive and degradation inhibitions, *α, κ*_*u*_ and *γ* are respectively increased to demonstrate their effect on reducing the stationary output (**X**_**2**_) variance. The plot compactly illustrates the variance-reduction effect of all three inhibition mechanisms using stochastic simulations (with 10^5^ trajectories for each parameter value) and using an approximate analytic formula that is derived based on a tailored moment-closure technique [16], [17]. The common parameters are *γ*_1_ = *γ*_2_ = *k*_1_ = 5, *k* = 3, *µ* = 10, *θ* = 2, *η* = 100, *δ* = 10. For additive inhibition, *κ*_*u*_ = 1 while *α ∈* [0, 25] is tuned. For competitive inhibition, 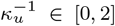 is tuned. Finally, for degradation inhibition, *κ*_1_ = 1 and *γ ∈* [0, 40] is tuned.

## VI. Comparison with previous antithetic Proportional-Integral Controllers

Previous antithetic proportional-integral controllers [16], [17], [19] are realized by invoking a direct negative feedback from the output species, unlike the design we presented here where the sequestration complex formed between **Z**_**1**_ and **Z**_**2**_ performs the negative feedback instead of the output. The key difference between these two approaches is that we obtain a filtered proportional component in the current design, while in previous designs we obtain a non-filtered proportional component. In this section, we demonstrate using, once again, the birth-death process as a plant, that the filtered PI gives rise to an additional degree of freedom that improves the flexibility in shaping the dynamic response. The demonstration is carried out once again using the pole placement technique performed in Section IV. By doing the same analysis for the non-filtered PI controller, the constraints on the achievable poles *s* = − *a* on the real axis can be written as

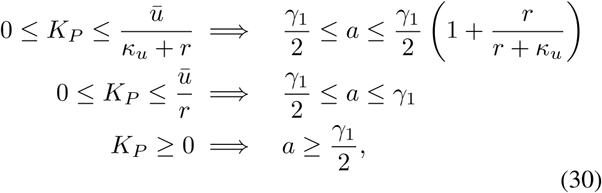

for the additive, competitive and degradation inhibitions, respectively, with *K*_*I*_ ≥ 0. Comparing equations (30) with (29), we conclude that using degradation inhibition, the same performance with filtered and non-filtered PI controllers can be achieved since we can place the poles arbitrarily far to the left in the complex plane in both cases. However, interestingly, for the other inhibition mechanisms the filtered PI controller where the complex **Z**_**3**_ acts as the inhibitor outperforms the previous PI controllers where the output **X**_**L**_ acts as the inhibitor.

## VII. Experimental Demonstration using the Cyberloop

For experimental validation, we use a biomolecular controller prototyping framework, the Cyberloop, proposed in [26]. This framework facilitates interfacing an *in silico* stochastic simulation of biomolecular controllers with a real biological target circuit *in vivo* via light stimulation and fluorescence observation. In this framework, multiple cells, placed under a microscope-coupled optogenetic platform [27], are tracked across time with each cell having its independent associated controller simulation. One can rapidly characterize and benchmark a given biomolecular controller using this hybrid framework without needing to implement the controller circuit *in vivo*. In this study, we used a previously engineered target circuit with nascent RNA as output molecule in *Saccharomyces cerevisiae* cells [27], and interfaced it with a stochastic simulation of the proposed filtered PI biomolecular controller network as shown in Figure 6(a).

**Fig. 6.**
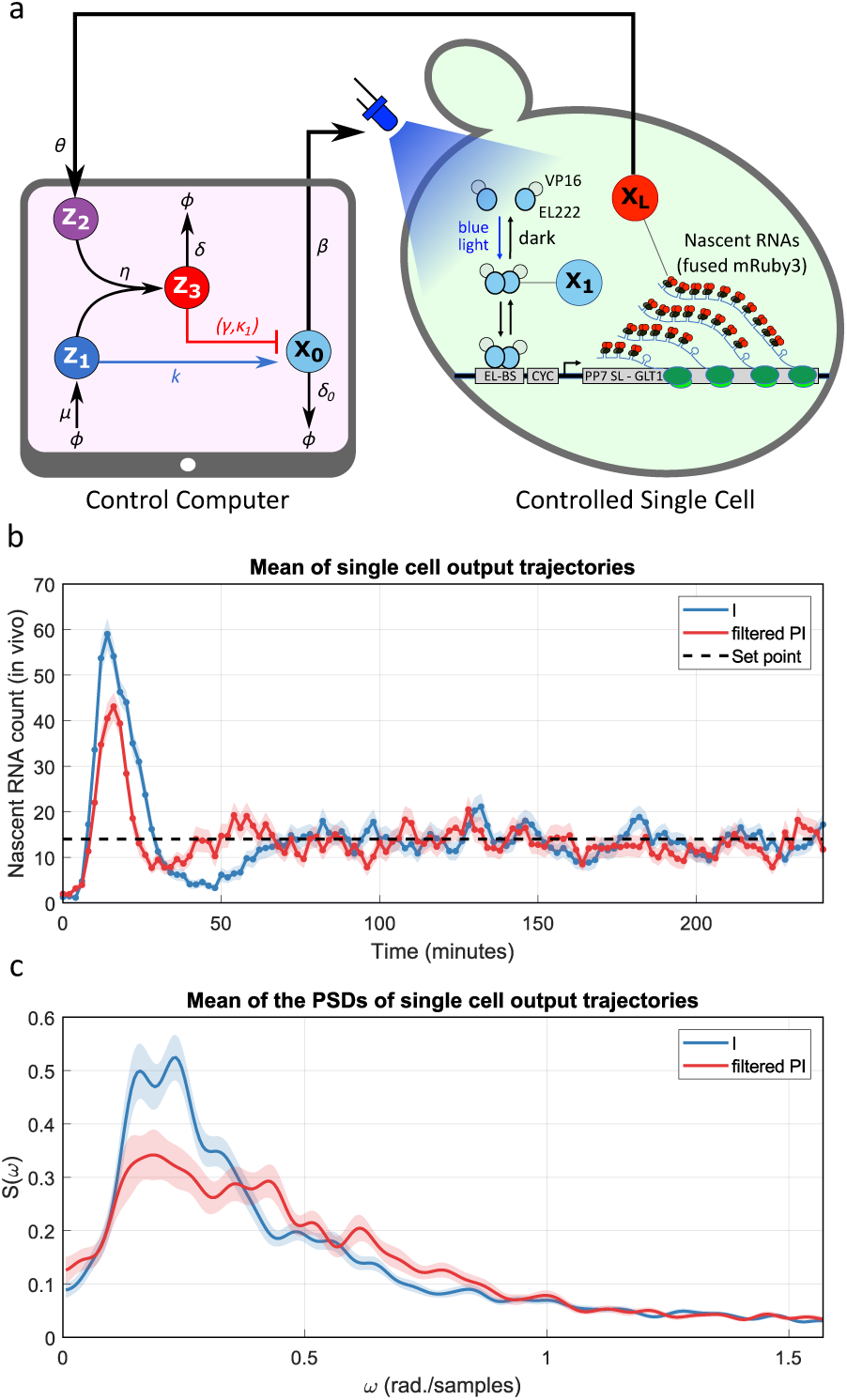
Cyberloop Experiment. (a) Implementation of the proposed filtered PI controller network on the Cyberloop platform. Following the controller prototyping strategy demonstrated in [26], the proposed filtered PI motif is simulated in stochastic setting on a control computer. The simulated controller is interfaced with a real biological circuit (target plant) genetically engineered in *Saccharomyces cerevisiae* cells proliferating under a microscope via an optogenetic platform [27]. This microscope-coupled platform allows one to stimulate single cells individually with light and observe/measure the fluorescence output in the cells in a parallel fashion. Here, the engineered target genetic circuit has nascent RNAs as output molecule, **X**_**L**_, which can be induced with blue light [27]. The output nascent RNAs can fuse with an available fluorescent protein (mRuby3) in the cell, and thus can be visualized and quantified from fluorescence microscopy images. In every imaging period, the quantified output values from multiple cells present in the microscopy field of view are fed to respective controller simulations (one controller per cell). These simulations compute input blue light intensities that the respective cells should be stimulated with in order to achieve the required output set-point tracking. Cells are then exposed to these input light intensities. This loop of fluorescence observation, controller simulation, and light stimulation are performed periodically with 2 minutes interval giving us a dynamic overview of the controller performance. Here, **Z**_**1**_, **Z**_**2**_ and **Z**_**3**_ are our proposed controller species (as mentioned previously) and **X**_**0**_ acts as a virtual plant species whose abundance defines the input light intensity for the cell. (b) Temporal response. This plot shows the average time trajectory of the output molecule from all target cells in two separate experiments with antithetic I and filtered PI controller motifs. The shaded region shows mean *±* standard error. (c) Frequency response. This plot shows the mean power spectral density (PSD) of output responses at steady-state in the two experiments. The shaded region shows mean *±* standard error. Experimental parameters: Number of tracked cells - 106 (I), 85 (filtered PI); *β* = 0.01, *θ* = 2, *µ* = 14 *× θ, η* = 5, *δ* = 1, *δ*_0_ = 0.5, *k* = 0.1, *γ* = 0 (for I) or 10 (for filtered PI), *κ*_1_ = 0.1, *α* = 1 [All units are in min^−1^].

We performed two separate experiments, one with antithetic integral (I) controller and another with our proposed filtered PI (with degradation inhibition) controller motif, and the experimental results are shown in Figure 6(b,c). As can be seen from the mean output time-trajectory plot in Figure 6(b), the filtered PI controller exhibits better transient dynamics with reduced overshoot and settling-time compared to that in the antithetic I case. Furthermore, the single-cell oscillations induced by the antithetic I motif for the given parameter values are also reduced in the filtered PI case. This is demonstrated in Figure 6(c), which shows the mean of the power spectral densities (PSDs) of all the tracked cell-output trajectories at steady state. Here, the filtered PI curve displays a lower peak compared to antithetic I case, indicating dampened stochastic single-cell oscillations.

## VIII. CONCLUSION

The biomolecular realization of controllers that ensure RPA is a fundamental requirement to stand against the uncertain and noisy nature of living cells. However, achieving RPA alone is often not enough, and pushing towards biomolecular controllers with high performance is essential. In this work, we have augmented the antithetic integral motif — a controller that ensures RPA — with an additional inhibition reaction to enhance the performance in both the deterministic and stochastic settings while maintaining the RPA property. One attractive feature of this controller is that it leverages a molecule which is already present, but inert, in the antithetic motif. We show that using this molecule as an inhibitor opens an extra dimension of design flexibility that is capable of shaping the transient dynamics and reducing the cell-to-cell variability.

## APPENDIX

### A. Pole Placement Calculations

In this section, we derive the bounds on the achievable poles in the three inhibition scenarios: degradation, competitive (multiplicative) and additive.

The supporting input to control the birth-death process can be calculated by setting the right hand side of the first equation in (4) to zero for *L* = 1 and *f* (*x*) = *γ*_1_*x*_*L*_ to obtain *ū* = *γ*_1_*r*, where *r* :=*µ/θ* is the desired set-point.

a. *Degradation Inhibition:* Substituting for (*K*_*P*_, *K*_*I*_, *ω*_0_) from (28) in (23) yields

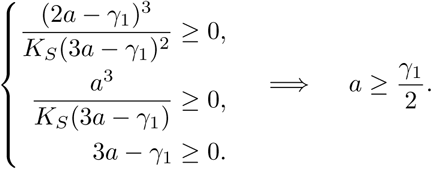
b. *Competitive Inhibition:* Substituting for (*K*_*P*_, *K*_*I*_, *ω*_0_) from (28) in (19) yields

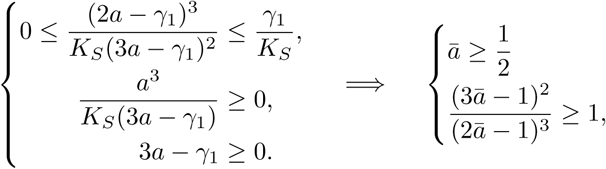

where *ā* := *a/γ*_1_. The second condition gives rise to a cubic polynomial given by 8*ā*^3^ *−* 21*ā*^2^ + 12*ā−* 2 *≤* 0, which can be solved to obtain *ā* ≤ *s*_*c*_ or *a* ≤ *a* ≤ *s*_*c*_*γ*_1_ where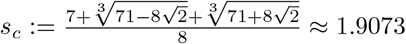.
c. *Additive Inhibition:* Substituting for (*K*_*P*_, *K*_*I*_, *ω*_0_) from (28) in (15) yields

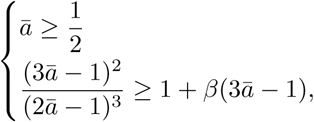

where *ā* := *a/γ*_1_ and 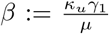. The second condition gives rise to a quartic polynomial that depends on the parameter group *β* and thus it is impractical to write down a closed formula for the solution of *ā* in terms of *β*. However, we will use a graphical argument to provide a bound for the solution. Define the following functions

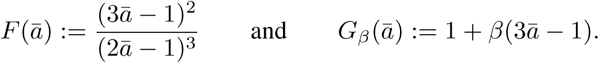

The intersection of these two functions is the solution of the quartic polynomial denoted by *s*_*a*_(*β*). Clearly, we have *s*_*a*_(0) = *s*_*c*_. Sketching the two functions on the same plot as shown in Figure 7 shows that 1*/*2 *≤ s*_*a*_(*β*) *≤ s*_*a*_(0) and *s*_*a*_(*β*) is a decreasing function of *β ≥* 0.

**Fig. 7.**
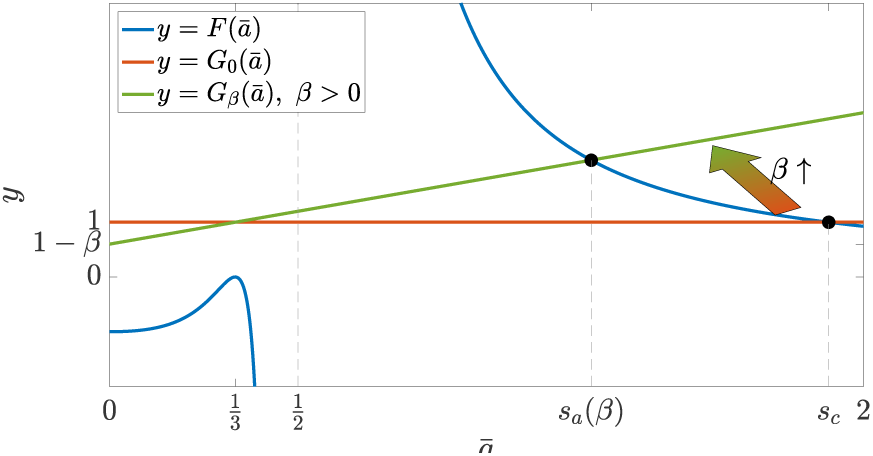
Bounds of the solution of the quartic polynomial.

### B. Moment Closure Approximation

In this section, we investigate the effect of the filtered PI controllers on the stationary (steady-state) behavior of the output species **X**_**L**_ in the stochastic setting. Particularly, we examine the stationary expectation 𝔼 _*π*_ [*X*_*L*_] and variance Var _*π*_ [*X*_*L*_].

#### 1) Stoichiometry Matrix & Propensity Function

Let *S* and *λ* respectively denote the stoichiometry matrix and propensity function of the general plant given in Figure 1(a). The stoichiometry matrix *S*_cl_ and propensity function *λ*_cl_ of the closed-loop system comprised of the plant given in Figure 1(a) controlled by the controller given in Figure 1(c) are given by

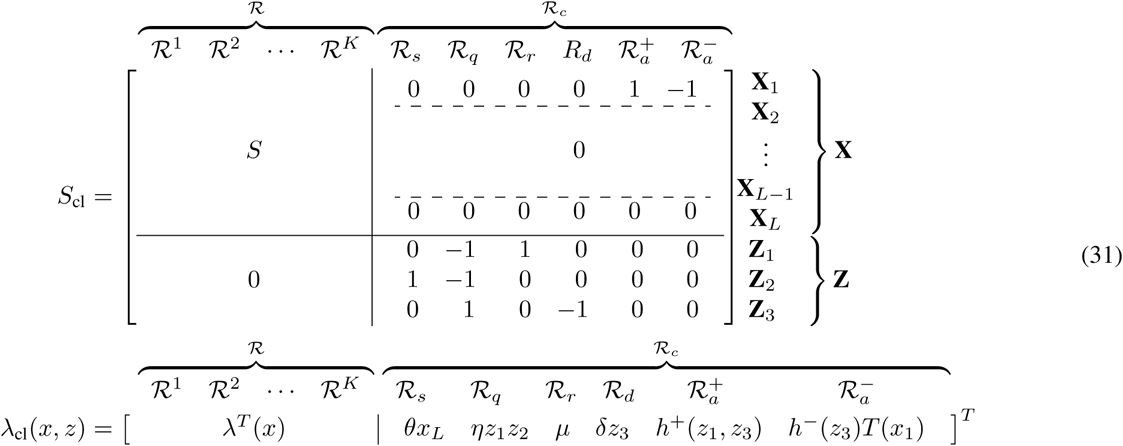

with *T* (*x*_1_) := *x*_1_*/*(*x*_1_ + *κ*_1_), and the negative actuation *h*^*−*^ is assumed to be influenced by *z*_3_ only.

#### 2) Stationary Expectation & RPA

The evolution of the expectations of the various species in the closed-loop network are simply given by the differential equation 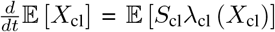, where *X*_cl_ := [*X*^*T*^ *Z*^*T*^]^*T*^. By substituting for the closed-loop stoichiometry matrix *S*_cl_ and propensity function *λ*_cl_, we obtain the following set of differential equations that describe the evolution of the expectations for an arbitraty plant.

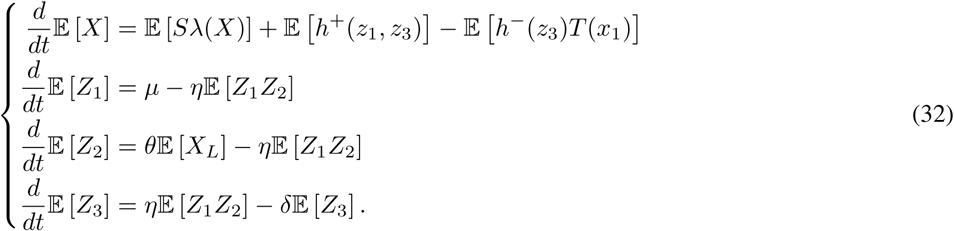

At stationarity, assuming that the closed-loop network is ergodic, the time derivatives converge to zero. Particularly, we have

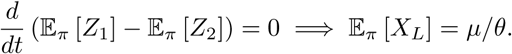

Thus RPA is ensured as long as the closed-loop network is ergodic.

#### 3) Stationary Variance Approximation

The goal of this section is to derive an approximate formula for the stationary variance of the output species **X**_**L**_ when the plant is controlled with the filtered PI controller of Figure 1(c). Unfortunately, a general analysis for an arbitrary plant cannot be done. As a case study, we consider the particular plant given in Figure 5 where the plant is a simple gene expression network, and it is connected in feedback with the filtered PI controller. Note that the analysis we performed can be generalized to any (affine-linear) plant with mono-molecular reactions. Even for this particular plant, one cannot derive an exact expression for Var_*π*_ [*X*_2_]. This is a consequence of the moment closure problem that stems from the inherent nonlinear nature of the antithetic motif (quadratic propensity: *ηz*_1_*z*_2_) and the nonlinear actuation propensities *h*^+^ and *h*^*−*^. However, a tailored moment-closure technique was proposed for the non-filtered PI case in [17] that exploits the fact that 𝔼_*π*_ [*Z*_1_*Z*_2_] =*µ/η≈* 0 for large *η*; and as a result assumes that *Z*_2_ remains close to zero. Furthermore, a linearized approximation of the actuation function *h* is also exploited to circumvent the moment closure problem. Extending this approximate technique to our filtered PI controller allows us to give a general (approximate) expression for Var_*π*_ [*X*_2_] that encompasses all three types of inhibitions. We now present the detailed mathematical derivations. First, we consider a general plant to write down the evolution equations of the variance. Then, we derive an approximate closed formula for the output stationary variance in the case of the particular (gene expression network) plant given in Figure 5.

Define the instantaneous covariance matrix of the closed-loop state variable as

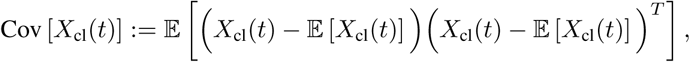

whose evolution is described by the following differential equation (we drop the time variable for notational convenience)

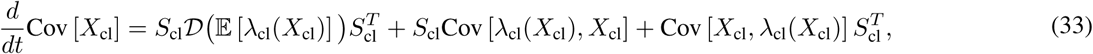

where *S*_cl_ and *λ*_cl_ denote the closed-loop stoichiometry matrix and propensity function, respectively. Note that *𝒟* is the diagonal operator such that for any vector *x, 𝒟*(*x*) is a diagonal matrix whose diagonal entries are *x*. Define the matrices and vector

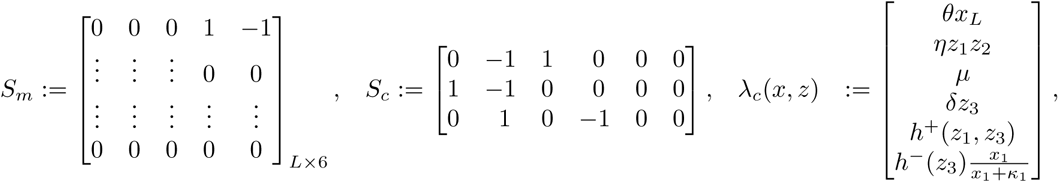

so that the closed-loop stoichiometry matrix and propensity function can be respectively written as

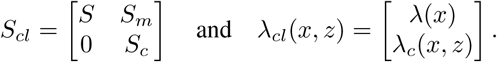

Recall that *h*^+^ and *h*^*−*^ are functions that can take particular forms depending on the adopted inhibition mechanism. By substituting the plant and controller components of *X*_cl_, *S*_cl_, and *λ*_cl_ in (33), we proceed as follows

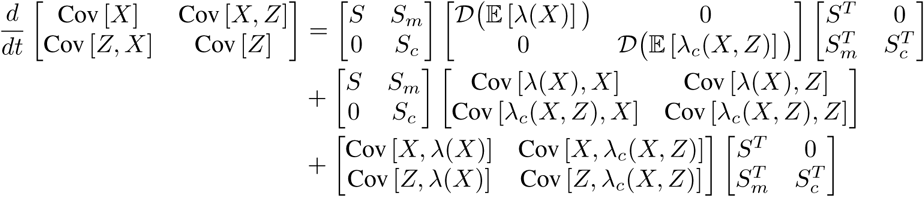

Thus we have

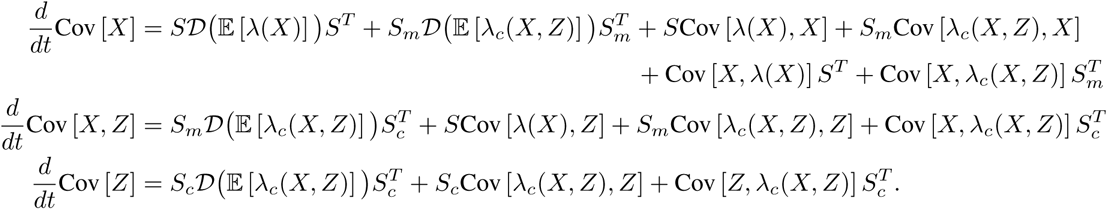

Next, by substituting for *S*_*m*_, *S*_*c*_ and *λ*_*c*_(*X, Z*) and doing some algebraic calculations, we obtain

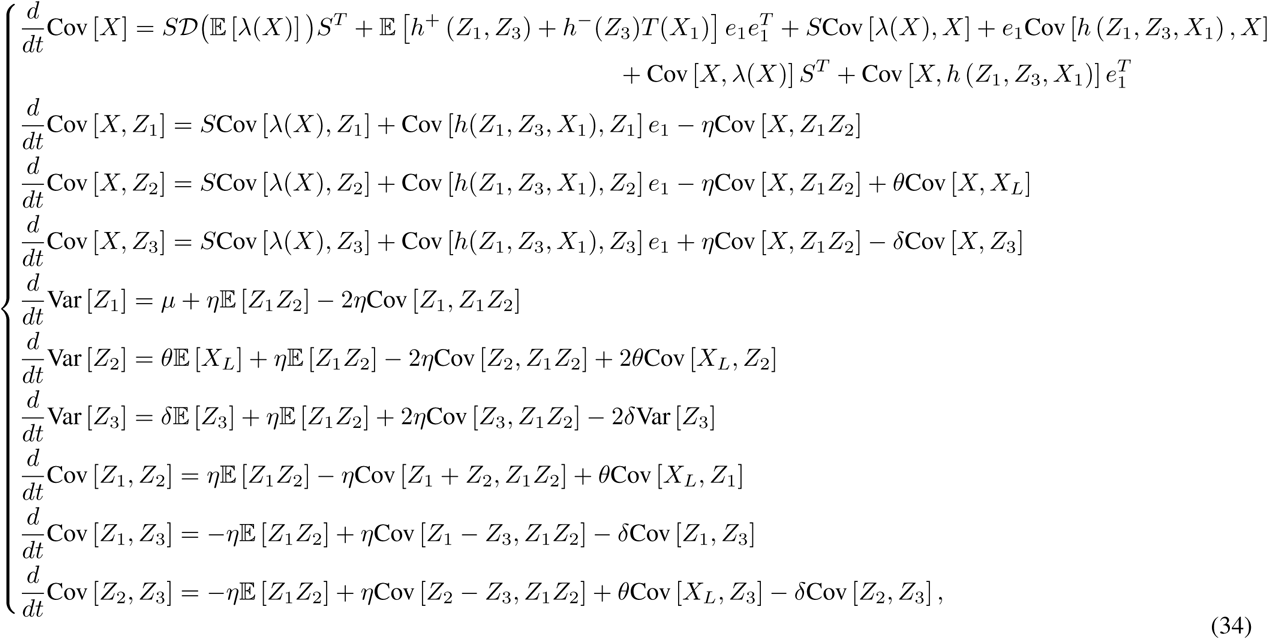

where the total actuation propensity function is defined as *h*(*z*_1_, *z*_3_, *x*_1_) := *h*^+^(*z*_1_, *z*_3_) − *h*^*−*^(*z*_3_)*T* (*x*_1_), and *e*_*i*_ is a vector whose entries are all zeros except the *i*^th^ entry is one. This set of differential equations describe the evolution of the various covariances in the closed-loop network. Observe that it does not involve first and second order moments only (expectations and covariances), but also third order moments like Cov [*X, Z*_1_*Z*_2_] and Cov [*Z*_1_ + *Z*_2_, *Z*_1_*Z*_2_] that have their own differential equations. This, in addition to the nonlinearity of *λ* and *h* (in general), give rise to the moment closure problem.

##### General Steady-State (Stationary) Analysis

Let 𝔼_*π*_ [·], Var_*π*_ [·], and Cov_*π*_ [·,·] denote the the stationary expectation, variance and covariance, respectively. Assuming that the closed-loop system is ergodic, the various time derivatives in (32) and (34) converge to zero at stationarity. Particularly, we have the following relationships that hold regardless of what the plant is

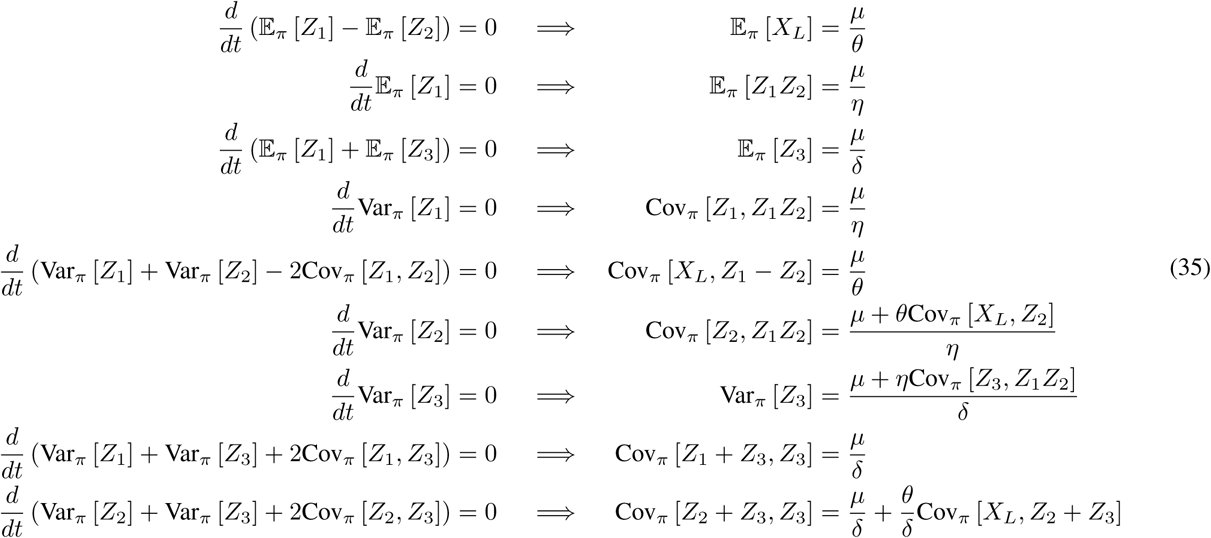

These relationships will be useful in what follows, particularly in the asymptotic limit as *η → ∞*.

##### Application to the Gene Expression Network

Consider the case where the plant is the gene expression network described in Figure 5. The plant stoichiometry matrix and propensity vector are given by

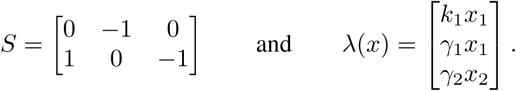

Then, by substituting *S* and *λ* in (32) and (34), we obtain the the following set of differential equations for the expectations and covariances

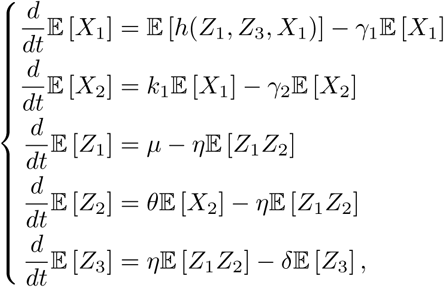

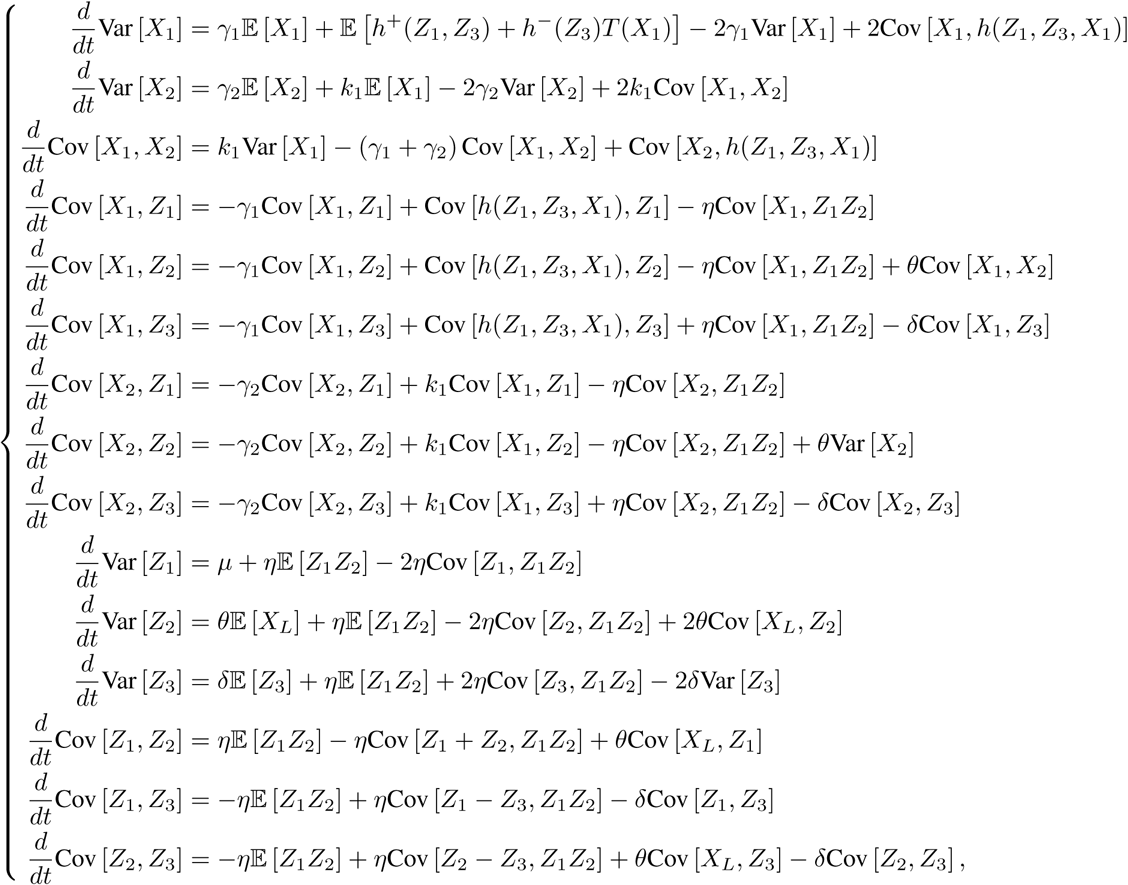

Assuming that the closed-loop system is ergodic, the time derivatives at stationarity are set to zero. We have

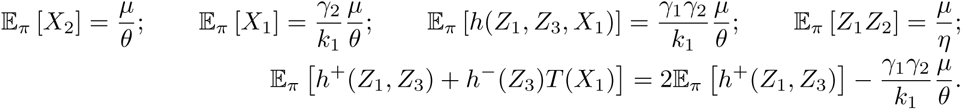

To compute the steady-state variance Var_*π*_ [*X*_2_], we use the following set of algebraic equations

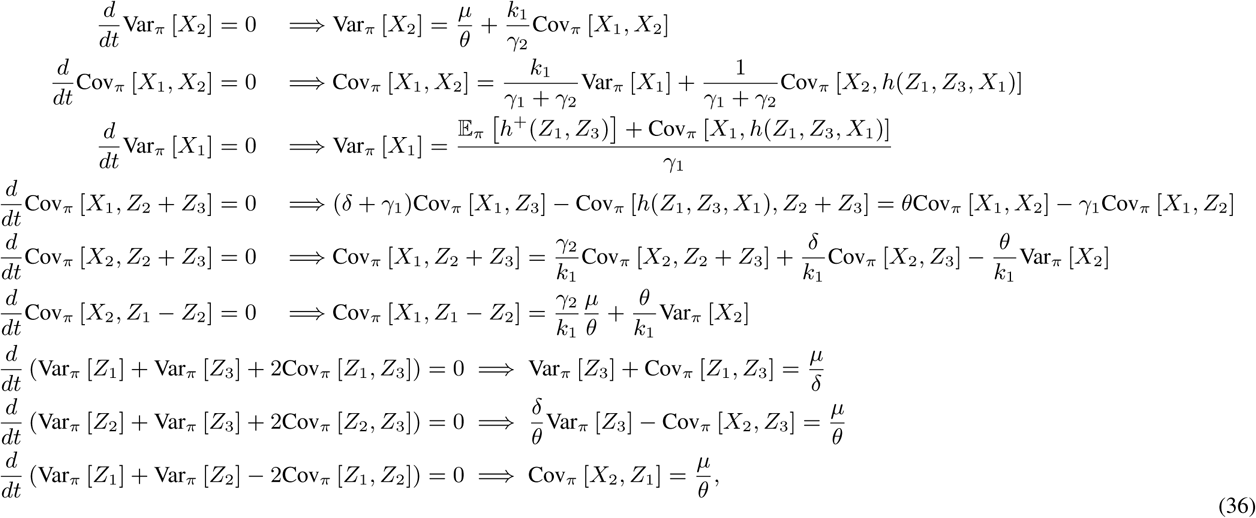

where the sixth equality follows by exploiting the fact that Cov_*π*_ [*X*_2_, *Z*_1_ − *Z*_2_] =*µ/θ* from (35). Observe that these algebraic equations cannot be solved exactly for Var_*π*_ [*X*_2_] because of the moment closure problem. However, to proceed, we give an approximation for the covariance terms Cov_*π*_ [*X*_*i*_, *h*(*Z*_1_, *Z*_3_, *X*_1_)] for *i* = 1, 2. The approximation essentially (1) exploits a second order Taylor expansion of the function *h* around the stationary expected values, and (2) exploits the fact that for large *η*, 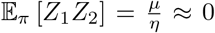 and *Z*_2_ is assumed to be close to zero unlike *Z*_1_ which takes positive values actuating the plant. In fact, these approximations are summarized below.

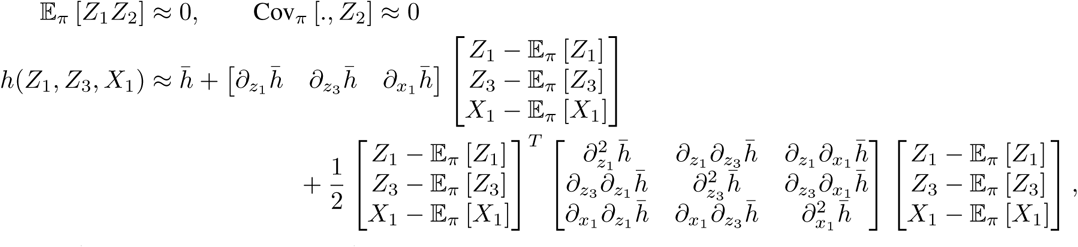

where 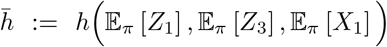. Resorting to [16, Supplementary Information Section S13] (with *X* := [*Z*_1_ *Z*_3_ *X*_1_]^*T*^, *F* (*X*) = *X*_1_, and *G*(*X*) = *h*(*Z*_1_, *Z*_3_, *X*_1_)), we can approximate Cov_*π*_ [*X*_1_, *h*(*Z*_1_, *Z*_3_, *X*_1_)] up to first order (or second order if the stationary distribution is close to a normal distribution) as

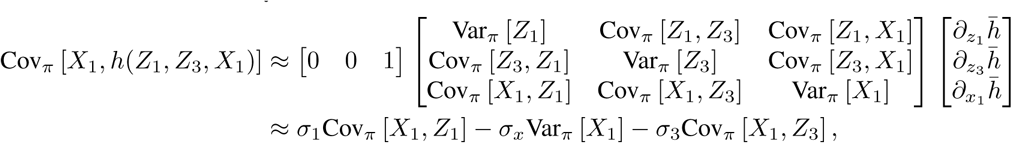

where

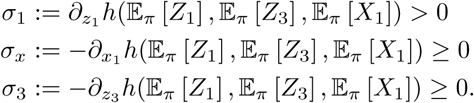

Similarly, we can approximate Cov_*π*_ [*X*_2_, *h*(*Z*_1_, *Z*_3_, *X*_1_)] as

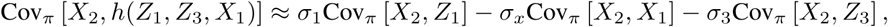

and

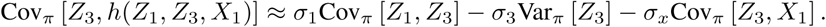

Invoking the approximations Cov_*π*_ [., *Z*_2_] *≈* 0 and a first order approximation for 𝔼_*π*_ [*h*^+^(*Z*_1_, *Z*_3_)], (36) can be approximated as a set of linear equations given by

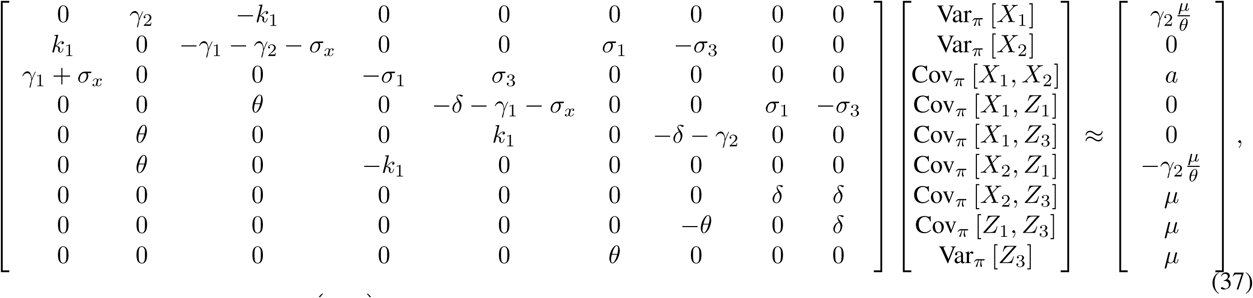

where 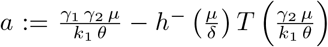. Finally using Cramer’s rule, one can obtain a closed approximate formula for the variance Var_*π*_ [*X*_2_] of the regulated output. The formula is computed symbolically in MATLAB, and is not shown here. Note that the three different inhibition scenarios can be computed using the following table.

**TABLE 1.**
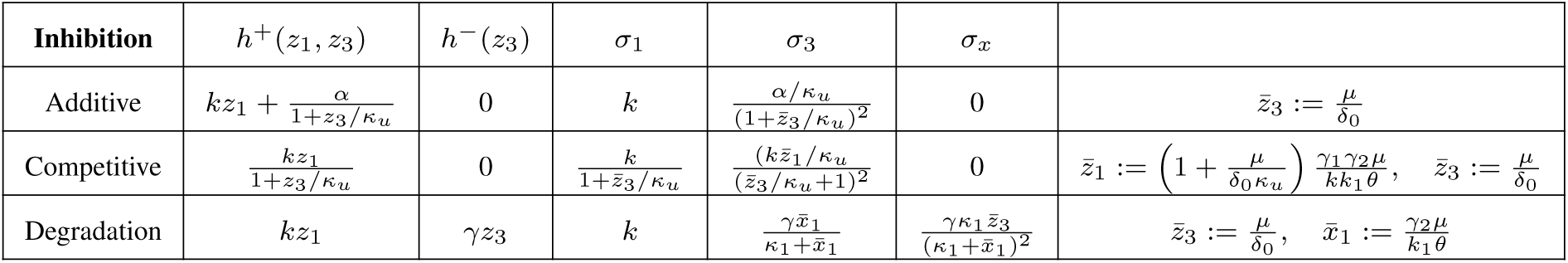
Actuation functions and partial derivatives for the three inhibition mechanisms.

